# Strain and lineage-level methylome heterogeneity in the multi-drug resistant pathogenic *Escherichia coli* ST101 clone

**DOI:** 10.1101/2020.06.07.138552

**Authors:** Melinda M. Ashcroft, Brian M. Forde, Minh-Duy Phan, Kate M. Peters, Leah W. Roberts, Kok-Gan Chan, Teik Min Chong, Wai-Fong Yin, David L. Paterson, Timothy R. Walsh, Mark A. Schembri, Scott A. Beatson

**Affiliations:** School of Chemistry and Molecular Biosciences, The University of Queensland, St Lucia, QLD, Australia; Australian Infectious Diseases Research Centre, The University of Queensland, St Lucia, QLD, Australia; Australian Centre for Ecogenomics, The University of Queensland, Brisbane, QLD, Australia; Division of Genetics and Molecular Biology, Institute of Biological Sciences, Faculty of Science, University of Malaya, Kuala Lumpur, Malaysia; International Genome Centre, Jiangsu University, Zhenjiang, China; UQ Centre for Clinical Research, The University of Queensland, Herston, QLD, Australia; Department of Medical Microbiology and Infectious Disease, Cardiff University, Cardiff, United Kingdom

**Keywords:** DNA methylation, Restriction-Modification Systems, Pacific Biosciences, Mobile Genetic Elements, epigenome

## Abstract

*Escherichia coli* Sequence Type (ST)101 is an emerging, multi-drug resistant lineage associated with carbapenem resistance. We recently completed a comprehensive genomics study on mobile genetic elements (MGEs) and their role in *bla*_NDM-1_ dissemination within the ST101 lineage. DNA methyltransferases (MTases) are also frequently associated with MGEs, with DNA methylation guiding numerous biological processes including genomic defence against foreign DNA and regulation of gene expression. The availability of Pacific Biosciences Single Molecule Real Time Sequencing data for seven ST101 strains enabled us to investigate the role of DNA methylation on a genome-wide scale (methylome). We defined the methylome of two complete (MS6192 and MS6193) and five draft (MS6194, MS6201, MS6203, MS6204, MS6207) ST101 genomes. Our analysis identified 14 putative MTases and eight N6-methyladenine DNA recognition sites, with one site that has not been described previously. Furthermore, we identified a Type I MTase encoded within a Transposon 7-like Transposon and show its acquisition leads to differences in the methylome between two almost identical isolates. Genomic comparisons with 13 previously published ST101 draft genomes identified variations in MTase distribution, consistent with MGE differences between genomes, highlighting the diversity of active MTases within strains of a single *E. coli* lineage. It is well established that MGEs can contribute to the evolution of *E. coli* due to their virulence and resistance gene repertoires. This study emphasises the potential for mobile genetic elements to also enable highly similar bacterial strains to rapidly acquire genome-wide functional differences via changes to the methylome.

**Impact Statement:** *Escherichia coli* ST101 is an emerging human pathogen frequently associated with carbapenem resistance. *E. coli* ST101 strains carry numerous mobile genetic elements that encode virulence determinants, antimicrobial resistance, and DNA methyltransferases (MTases). In this study we provide the first comprehensive analysis of the genome-wide complement of DNA methylation (methylome) in seven *E. coli* ST101 genomes. We identified a Transposon carrying a Type I restriction modification system that may lead to functional differences between two almost identical genomes and showed how small recombination events at a single genomic region can lead to global methylome changes across the lineage. We also showed that the distribution of MTases throughout the ST101 lineage was consistent with the presence or absence of mobile genetic elements on which they are encoded. This study shows the diversity of MTases within a single bacterial lineage and shows how strain and lineage-specific methylomes may drive host adaptation.

**Data Summary:** Sequence data including reads, assemblies and motif summaries have previously been submitted to the National Center for Biotechnology Information (https://www.ncbi.nlm.nih.gov) under the BioProject Accessions: PRJNA580334, PRJNA580336, PRJNA580337, PRJNA580338, PRJNA580339, PRJNA580341 and PRJNA580340 for MS6192, MS6193, MS6194, MS6201, MS6203, MS6204 and MS6207 respectively. All supporting data, code, accessions, and protocols have been provided within the article or through supplementary data files.

## Introduction

*Escherichia coli* sequence type (ST)101 is a pathogenic clone that has recently been associated with urinary tract and bloodstream infections in humans [1-4]. ST101 represents one of the major, emerging *E. coli* clones associated with the carriage of the *bla*_NDM-1_ gene, causing carbapenem resistance [1, 5-9]. Recently, we undertook the most comprehensive genomics study on mobile genetic elements (MGEs) and their role in *bla*_NDM-1_ dissemination within the ST101 lineage to date [10]. We sequenced the genomes of seven *bla*_NDM-1_-positive ST101 isolates using Pacific Biosciences (PacBio) Single Molecule Real Time (SMRT) sequencing, generating two complete (MS6192 and MS6193) and five high-quality draft genomes (MS6194, MS6201, MS6203, MS6204, MS6207) [10]. Using an additional thirteen previously published and publicly available draft ST101 genomes, we showed that ST101 strains formed two distinct clades (Clades 1 and 2) with clustering based on infection site, *fimH* profile and antimicrobial resistance gene repertoire. Notably multidrug resistance and the carriage of the *bla*_NDM-1_ gene were restricted to a subset of Clade 1 isolates. ST101 strains have a variable mobile genetic element (MGE) complement including prophages, genomic islands, transposons, and plasmids that encode genes for virulence, fitness, and antimicrobial resistance. Many MGEs also contained DNA methyltransferase (MTase) genes, which may result in differential methylation patterns.

In bacteria, DNA methylation is catalysed by MTases, where it guides many biological processes including defence mechanisms against foreign DNA, DNA replication and repair, timing of transposition and regulation of gene expression [11]. Three methylated nucleotides are known to occur in bacteria: N6-methyladenine, (^6m^A), N4-methylcytosine (^4m^C) and C5-methylcytosine (^5m^C) [12]. MTases are often encoded alongside, or as part of, restriction endonucleases (REases), which have the same DNA recognition site, forming restriction-modification (RM) systems that play a central role in defence against foreign, invading DNA [13]. Additionally, MTases can act independently of REases and such DNA-modifying enzymes are known as orphan MTases. MTases and RM systems are ubiquitous and extremely diverse in prokaryotes, and are classified into four major groups: Type I, II, III and IV based on subunit composition, DNA recognition site specificity, site of cleavage and reaction substrates (for a comprehensive review: [14]). In *E. coli*, RM systems and orphan MTases are most commonly Type I or Type II [14]. Type I systems are comprised of three subunits: restriction (R), modification (M) and specificity (S) [15]. The S subunit contains the DNA target recognition domain (TRD) and recognises bipartite asymmetric recognition sequences separated by 4-9 degenerate bases [15]. Type II systems are the most widespread, and in their simplest form comprise separate R and M genes with identical DNA binding specificity, and often recognise 4-6 bp palindromic sequences [16]. Exceptions include Type IIG, where the R and M subunits are contained in one polypeptide and in general, bind to short, non-palindromic sequences, resulting in hemi-methylation [17, 18]. Knowledge of MTase binding specificities is critical for pairing motifs with their cognate MTase.

Genes encoding DNA MTases have been identified in most prokaryote genomes available to date [13, 19]. However, despite the rapid growth of genomic information in public databases, epigenomic information such as methylation has lagged due to methodological limitations of previous technologies [20]. PacBio SMRT sequencing produces long reads, enabling the resolution of complex genetic structures such as MGEs and *de novo* assembly of complete bacterial chromosomes and plasmids [21]. Additionally, SMRT sequencing can be used to identify DNA modifications such as methylation at a single base resolution, based on the kinetics of the sequencing reaction [20]. PacBio SMRT sequencing can directly detect ^6m^A and ^4m^C modifications due to their robust kinetic signatures, however it is only moderately sensitive for ^5m^C modifications [22]. The impact of SMRT sequencing on cataloguing genome-wide methylation in bacteria has been demonstrated recently, with the complete methylome of hundreds of bacterial pathogens and environmental species now characterised (for example [23]). MTases and RM systems that have been characterised in bacteria are often encoded on MGEs [13, 19] and have additional biological roles including the generation of genomic diversity required for host fitness [13, 24]. However, there are relatively few studies on the genomic context and functional and evolutionary consequences of most identified MTases.

Except for Ashcroft *et al*. [10], there have been limited genomics studies of the *E. coli* ST101 lineage and no methylome analyses to date. Here, we present the first methylome analysis of ST101 using PacBio SMRT sequencing data for seven *E. coli* ST101 genomes. We defined the patterns of DNA methylation across all seven ST101 genomes, pairing recognition sites with their cognate MTase. Notably, we found a functional Type I RM system encoded within a Transposon 7-like Transposon (Tn) was responsible for extensive methylome differences in otherwise identical strains. By including an additional 13 previously published, draft ST101 genomes, we found that the majority of MTases were encoded on variably distributed MGEs, giving the potential for an unprecedented level of differential methylation within a single *E. coli* lineage.

## Methods

### SMRT sequencing and whole-genome detection of methylated bases

Genomic DNA (gDNA) of seven *E. coli* ST101 strains; MS6192, MS6193, MS6194, MS6201, MS6203, MS6204 and MS6207 was extracted from overnight cultures and sequenced on either a PacBio RSI or RSII sequencer as previously described [10]. Detection of methylated bases and the identification of associated methyltransferase recognition sites across the seven genomes (2 complete, 5 draft), was performed using the RS_Modification_and_Motif_Analysis protocol within the SMRT Analysis suite v2.3.0. Interpulse durations (IPDs) were calculated based on the kinetics of the nucleotide incorporation and were processed as previously described [20]. Sequence motifs were identified using Motif Finder v1, implemented in the SMRT Portal v2.3.0. Quality value cut-offs of 20 and 30 were applied for the draft and complete genomes, respectively. Here we report only ^6m^A methylation. As the DNA was not Ten-eleven translocation (Tet) treated prior to sequencing, ^5m^C modifications were not quantitated and ^4m^C modifications were not identified in any genome.

### Analysis of methyltransferase target site enrichment in gene regulatory regions

Putative gene regulatory (promoter) regions were defined as up to 300 bp upstream of the start codon of each CDS. To identify RM.EcoST101V recognition sites that were within intergenic regions we used the Bedtools v2.23.0 [25] closest flag, which reports the nearest genomic distance between recognition sites and CDSs. Sites that were within or overlapped the ends of CDSs were removed. A list of all protein-coding genes that contained a 5’-^6m^**A**CGN_5_G<u>T</u>TG-3’ site within 300 bp upstream of a start codon in MS6193 was generated (Supplementary Dataset, Table S1). This was used as a target gene list to compare with a background gene list formed by all genes within the *E. coli* K12 genome. If a gene was within an operon, all members of the operon were included. This target and background gene list comparison was performed using the functional enrichment analysis within the Database for Annotation, Visualization and Integrated Discovery (DAVID) v6.7 [26, 27]. Genes were annotated based on known function, gene ontologies and pathways. Results were determined as significant if post-hoc Benjamini-Hochberg correction for multiple testing reported a P value of <0.05.

### Methyltransferase diversity

To further analyse the distribution of MTases across the *E. coli* ST101 lineage, 13 additional published and publicly available ST101 draft genomes were downloaded from Genbank or the SRA as previously described [10]. An additional eight ExPEC complete genomes (accession details available in Supplementary Dataset, Table S2) were also included to emphasise ST101 lineage specific MTases. Active MTase genes identified in the *E. coli* ST101 draft genomes, all MTase genes from MS6192 and MS6193 and MTases from the REBASE Gold Standard database were searched against the 20 ST101 genomes and eight ExPEC complete genomes (Nucleotide Blast, ≥90% nucleotide identity and >95% sequence coverage) with redundancy removed. The presence or absence of MTase genes were visualised using Seqfindr (http://github.com/mscook/seqfindr).

## Results

### *E. coli* ST101 complete genomes MS6192 and MS6193 encode an almost identical complement of DNA methyltransferases

To characterise the role of methylation in shaping the *E. coli* ST101 lineage, we first defined the MTase complement of two near-identical Clade 1 *E. coli* ST101 strains (MS6192 and MS6193) for which we had previously determined the complete genomes [10]. The MS6192 genome encodes 12 putative MTases, with 8 on the chromosome, two on the large *bla*_NDM-1_-positive plasmid pMS6192A-NDM and one on each of the other large plasmids (pMS6192B and pMS6192C) (Table 1). Three chromosomal MTases correspond to enzymes that have been characterised in other *E. coli* strains, including Dam (5’-G^6m^**A**<u>T</u>C-3’), Dcm (5’-**C**^5m^CW<u>G</u>G-3’) and a homolog of the orphan MTase gene *yhdJ* encoding M.EcoST101III (5’-A<u>T</u>GC^6m^**A**T-3’) (by convention, underlined bases indicate methylation on the opposite strand), which has previously been reported to be inactive in other *E. coli* [28]. Additionally, we identified two Dam-like, orphan, Type II MTases (M.EcoST101I and M.EcoST101II) of unknown specificity located on the prophages Phi2 and Phi6, respectively. Three orphan, Type II MTases with unknown recognition sites also exist, with M.EcoST101VI and M.EcoST101VII encoded on Phi7 and M.EcoST101VIII encoded on the *bla*_NDM-1_-positive F-type plasmid pMS6192A-NDM. Also present are two orphan, Type II MTases encoded on each of the plasmids pMS6192B (M.EcoST101X) and pMS6192C (M.EcoST101XI); the recognition sites of these two MTases remains unknown. The remaining two MTases correspond to Type I RM systems. RM.EcoST101V is carried on the chromosome in an ST101 region of difference (RD12), with RM.EcoST101IX encoded on pMS6192A-NDM.

**Table 1.**
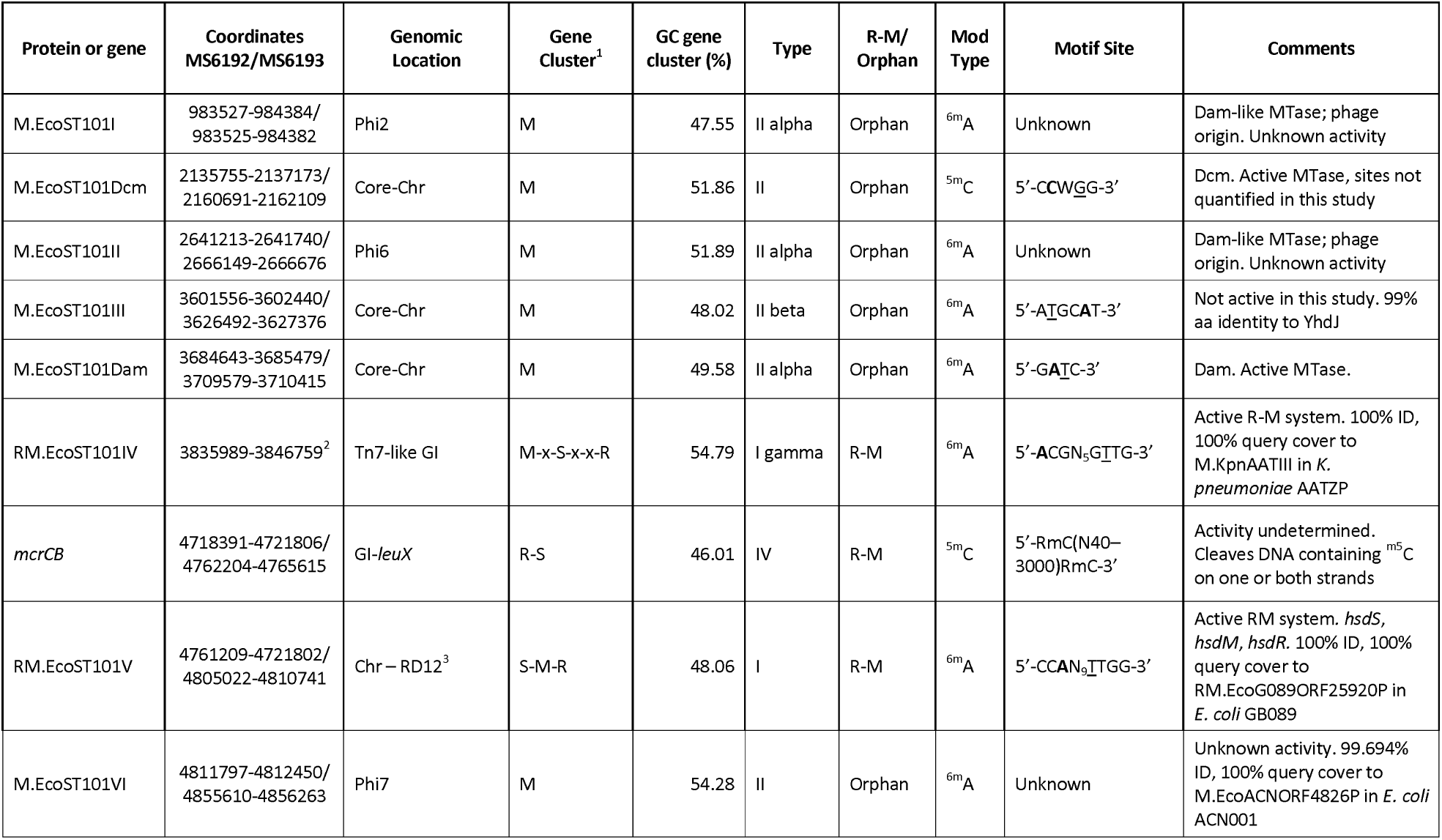
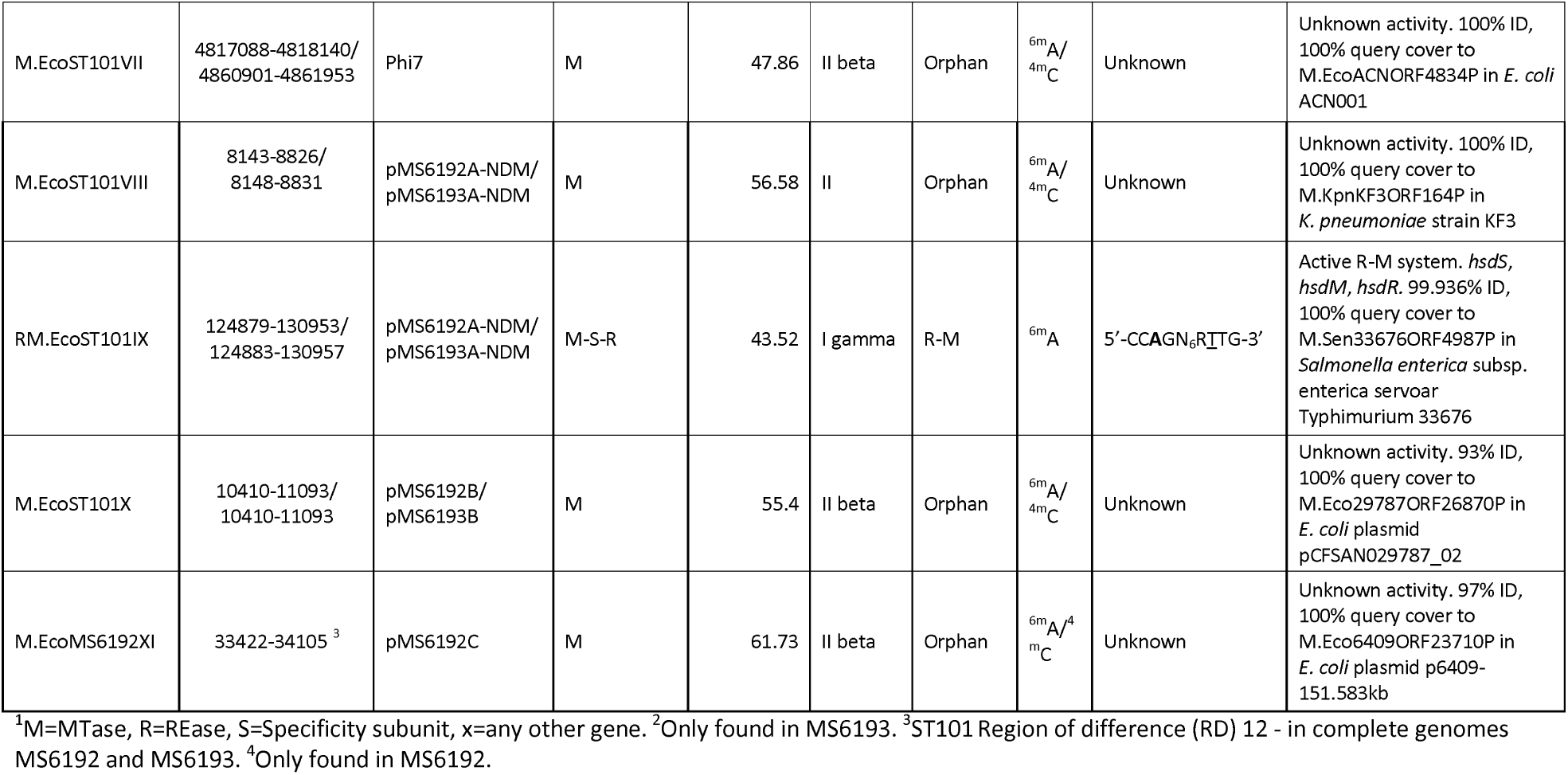
Summary of DNA methyltransferases and Restriction-Modification systems identified in the *E. coli* ST101 complete genomes MS6192 and MS6193.

Consistent with their close evolutionary relationship, MS6193 encodes all MTases found in MS6192 except for the Type II MTase M.EcoST101XI, as there is no plasmid corresponding to pMS6192C in the MS6193 genome. MS6193 also encodes an additional Type I RM system RM.EcoST101IV, carried on the Tn7-like Transposon that is not found in the MS6192 genome. Despite MS6193 also encoding an additional prophage (Phi8), no MTases were identified on this MGE.

### *E. coli* ST101 MS6192 and MS6193 genomes are differentially methylated

The genome-wide distribution of methylated bases in the complete genomes of MS6192 and MS6193 was determined using PacBio SMRT sequencing. Three distinct MTase recognition sites were detected as ^6m^A methylated in both genomes: 5’-G^6m^**A**<u>T</u>C-3’, 5’-CC^6m^**A**N_9_<u>T</u>TGG-3’ and 5’-CC^6m^**A**GN_6_R<u>T</u>TG-3’. The recognition site 5’-^6m^**A**CGN_5_G<u>T</u>TG-3’ was also detected in MS6193, but not MS6192 (Figure 1). To assign methylated sites to their cognate enzyme we used a process of elimination. As expected, one of the four recognition sites matched the well-characterised orphan, Type II MTase Dam (M.EcoST101Dam), with known specificity: 5’-G^6m^**A**<u>T</u>C-3’. Recently, the Type I RM recognition site 5’-CC^6m^**A**N_9_<u>T</u>TGG-3’ was identified in the *E. coli* strain GB089, however a cognate MTase was not assigned in REBASE [14]. Nucleotide comparisons of all Type I RM systems in GB089 and MS6192 revealed a single match between RM.EcoG089ORF25920P and RM.EcoST101V (100% identity), thus we deduce that the 5’-CC^6m^**A**N_9_<u>T</u>TGG-3’ recognition site is methylated by RM.EcoST101V. To investigate the third recognition site shared by both MS6192 and MS6193 (5’-CC^6m^AGN_6_R<u>T</u>TG-3’), we searched the motif against REBASE [14] and confirmed that it matches the recognition site of RM.Eco067II, identified in the *E. coli* strain AR_0067 (Genbank accession: CP032258). This motif is characteristic of a Type I RM system and with only one other Type I RM system identified in MS6192, we deduce that RM.EcoST101IX is responsible for methylation of the 5’-CC^6m^**A**GN_6_ R<u>T</u>TG-3’ site. Amino acid comparisons of the specificities subunits (HsdS) S.Eco067II and S.EcoST101XI confirm this match (99.77% identity, single amino acid substitution Y204H). The final ^6m^A recognition site 5’-^6m^**A**CGN G<u>T</u>TG-3’, detected only in MS6193, has previously been identified in the *Klebsiella pneumoniae* strain AATZP [29], and was assigned to the Type I RM system RM.KpnAATIII in REBASE. A nucleotide comparison showed that RM.KpnAATIII and RM.EcoST101IV share 100% nucleotide identity. Thus the Type I RM system RM.EcoST101IV in MS6193 must be responsible for methylation at 5’-^6m^**A**CGN_5_G<u>T</u>TG-3’. Also observed in both genomes were variations of the 5’-C^5m^**C**W<u>G</u>G-3’ motif, which is characteristic of Dcm methylation. Despite the presence of M.EcoST101Dcm in both genomes, the DNA was not Ten-eleven translocation (Tet)-treated and the SMRT sequencing coverage is lower than 250X, therefore accurate detection and quantification of ^5m^C in these genomes was limited (Supplementary Dataset, Table S3).

**Figure 1.**
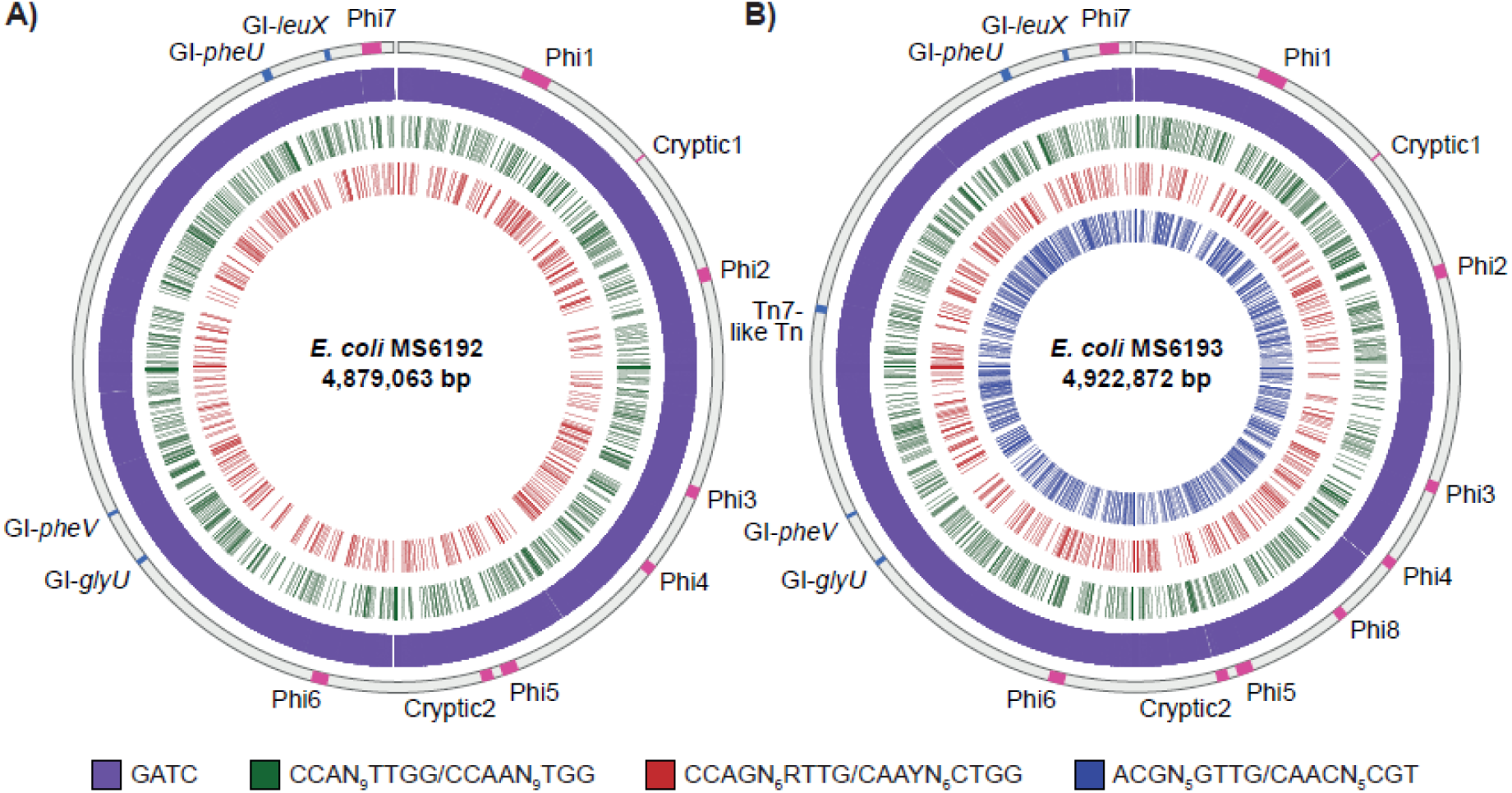
Circos plot displaying the distribution of ^m6^A methylated bases in the *E. coli* MS6192 and MS6193 chromosomes. The location of MGEs on the chromosome is indicated on the outermost track; Prophages (Pink), Genomic Islands and Transposons (Blue). The remaining tracks display the location of methylated recognition sites for each motif. A) For *E. coli* MS6192, from outer to inner: GATC, purple (M.EcoST101Dam); CCAN_9_TTGG/CCAAN_9_TGG, green (RM.EcoST101V); CCAGN_6_RTTG/CAAYN_6_CTGG, red (RM.EcoST101VIII). B) For *E. coli* MS6193, from outer to inner: GATC, purple (M.EcoST101Dam); CCAN_9_TTGG/CCAAN_9_TGG, green (RM.EcoST101V); CCAGN_6_RTTG/CAAYN_6_CTGG, red (RM.EcoST101VIII) and ACGN_5_GTTG/CAACN_5_CGT, blue (RM.EcoST101V).

#### RM.EcoST101IV may have acquired a secondary role in gene regulation

The additional 18.9 Kb Tn7-like Tn in MS6193 encoding RM.EcoST101IV (Supplementary Figure S1) is one of the major differences between the two complete genomes MS6192 and MS6193. We hypothesised that the acquisition of this additional RM system may lead to functional differences between the MS6192 and MS6193 strains. While the functional role of M.EcoST101IV is not currently known, the majority of the 788 5’-^6m^**A**CGN_5_G<u>T</u>TG-3’ sites (96%) are found in coding regions of the MS6193 genome. As methylation sites in intragenic regions are more likely to be associated with gene regulation [23], this suggests a primary role for RM.EcoST101IV in defence against foreign DNA. We also identified the presence of two methylated 5’-^6m^**A**CGN_5_G<u>T</u>TG-3’ sites within the Tn7-like Tn itself, found in MS6193_03822 encoding a putative DNA repair ATPase (UniProt), immediately upstream of the *hsdS* gene of RM.EcoST101IV. Although the functional consequence of these methylated sites is unknown, this may protect the Tn7-like Tn itself from degradation.

We identified 31 5’-^6m^**A**CGN_5_G<u>T</u>TG-3’ sites on the MS6193 chromosome and four on the plasmid pMS6193A-NDM that were in intergenic regions. Of these sites, all but one were within 300 bp of a start codon, which highlights the potential for RM.EcoST101IV to have acquired a secondary role in gene regulation (Supplementary Dataset, Table S1). From this, we generated a target gene list of 36 genes (including all genes within an operon if the RM.EcoST101IV site was within the putative promoter region for that operon). These genes include the transcriptional regulators *mcbR* and *fimZ, mntP* (putative manganese efflux pump), *yejO* (predicted autotransporter outer membrane protein), *ydcM* (putative transposase and virulence-associated protein) and *pagN* (outer membrane protein and virulence-associated protein). Notably, a single RM.EcoST101IV site was overlapping the start of an IS*26* element, which is 124 bp upstream of the truncated IS*Aba125* element that provides the −35 promoter region for the *bla*_NDM-1_ gene [30], responsible for carbapenem resistance in this lineage. Carbapenem Minimum Inhibitory Concentration (MIC) values were however identical between MS6192 and MS6193 except for Doripenem, which was lower in MS6192 by one serial dilution, yet still above the resistance breakpoint [10]. Despite no significant enrichment of functional pathways, these genes were primarily associated with cofactor binding, cell walls and membranes, ATP binding, nucleotide binding and metal ion binding (Supplementary Dataset, Table S4).

### Variation in *E. coli* ST101 Clade 1 methylomes is associated with variability in the accessory genome

To further investigate the ST101 methylome diversity, we included in our analyses five draft genome assemblies (MS6194, MS6201, MS6203, MS6204 and MS6207) that we have previously described [10]. In total, we identified four active ^6m^A MTases described above in MS6192 and MS6193, plus an active Type I RM system (RM.EcoST101XII), found only in MS6201 and MS6203 (Table 2). We also identified a novel ^6m^A Type II-like motif (5’-AGG^6m^**A**NTT-3’) in MS6203, resulting in hemi-methylation, however we could not definitively match it to its cognate MTase. The following cases illustrate several different scenarios that lead to differential methylation due to MGE-borne MTases in the ST101 lineage.

**Table 2.** Summary of DNA methyltransferase recognition sites identified in the PacBio sequenced *E. coli* ST101 strains in this study.

| Motif<sup>1</sup> | Name | Position of MTase | Mod Type | Mod Pos | Strains | No. sites in genome | No. sites detected | Methylated (%) | Mean QV | Mean IPD Ratio | Mean Site coverage |
| --- | --- | --- | --- | --- | --- | --- | --- | --- | --- | --- | --- |
| **GATC** | M.EcoST101Dam | Core-genome | <sup>6m</sup>A | 2 | MS6207 | 43154 | 38809 | 89.93 | 35.07 | 5.97 | 16.25 |
|  |  |  |  |  | MS6201 | 42732 | 39639 | 92.76 | 36.62 | 5.94 | 17.32 |
|  |  |  |  |  | MS6203 | 42144 | 39517 | 93.77 | 37.26 | 5.89 | 17.91 |
|  |  |  |  |  | MS6204 | 43182 | 37737 | 87.39 | 34.34 | 5.99 | 15.67 |
|  |  |  |  |  | MS6192 | 41748 | 41664 | 99.8 | 95.05 | 6.09 | 53.82 |
|  |  |  |  |  | MS6194 | 41956 | 39948 | 95.21 | 38.88 | 5.99 | 18.87 |
|  |  |  |  |  | MS6193 | 41646 | 41458 | 99.55 | 73.68 | 5.89 | 41.53 |
| **ACGN<sub>5</sub>GTTG/CAACN<sub>5</sub>CGT** | RM.EcoST101IV | Tn7-like GI | <sup>6m</sup>A | 1/3 | MS6193 | 788 | 783/777 | 99.37/98.6 | 69.53/68.31 | 6.71/6.36 | 41.22/41.26 |
| **CCAN<sub>9</sub>TTGG/CCAAN<sub>9</sub>TGG** | RM.EcoST101V | RD12<sup>2</sup> | <sup>6m</sup>A | 3/4 | MS6203 | 518 | 487/468 | 94.02/90.35 | 38.02/35.67 | 7.39/6.43 | 18.35/18.08 |
|  |  |  |  |  | MS6192 | 502 | 502/501 | 100/99.8 | 92.37/86 | 7.73/6.73 | 53.47/53.04 |
|  |  |  |  |  | MS6194 | 499 | 476/466 | 95.39/93.39 | 39.26/36.96 | 6.42/3.22 | 18.95/20.51 |
|  |  |  |  |  | MS6193 | 492 | 491/485 | 99.8/99.55 | 71.84/68.77 | 7.67/6.87 | 40.89/41.53 |
| **CCAGN<sub>6</sub>RTTG/CAAYN<sub>6</sub>CTGG** | RM.EcoST101IX | pIP1206-like FII plasmid | <sup>6m</sup>A | 3/3 | MS6204 | 853 | 756/698 | 88.63/81.83 | 35.05/32.33 | 8.79/6.1 | 15.67/15.43 |
|  |  |  |  |  | MS6192 | 821 | 821/815 | 100/99.27 | 95.12/86.69 | 8.99/5.97 | 53.72/52.39 |
|  |  |  |  |  | MS6194 | 833 | 807/765 | 96.88/91.84 | 39.3/36.16 | 8.78/5.91 | 18.62/18.57 |
|  |  |  |  |  | MS6193 | 820 | 819/808 | 99.88/95.54 | 73.75/69.92 | 9.06/5.95 | 40.99/40.55 |
| **GCAN<sub>5</sub>GTTC/GAACN<sub>5</sub>TGC** | RM.EcoST101XII | GI-*pheU* | <sup>6m</sup>A | 3/3 | MS6201 | 593 | 557/529 | 93.93/89.21 | 36.42/34.56 | 7.39/6.35 | 17.37/16.83 |
|  |  |  |  |  | MS6203 | 577 | 544/519 | 94.28/89.95 | 36.44/34.82 | 7.34/6.08 | 17.57/17.35 |
| **AGGANTT** | N/A | Unknown | <sup>6m</sup>A | 4 | MS6203 | 1986 | 1803 | 90.79 | 35.53 | 5.82 | 17.88 |

<sup>1</sup>By convention, bold bases and underlined bases indicate methylation on the forward and reverse strand, respectively. <sup>2</sup>ST101 Region of Difference (RD) 12 – defined in our complete genomes MS6192 and MS6193. QV = Quality Value

#### Differential methylation due to truncation of the DNA specificity gene in RM.EcoST101V

Methylation at the 5’-CC^6m^**A**N_9_T<u>T</u>GG-3’ site, which we have assigned to the Type I RM system RM.EcoST101V, is also present in two of the five draft genomes: MS6194 and MS6203. RM.EcoST101V is encoded within the 7.2 Kb ST101 region of difference 12 (RD12) of the complete genomes MS6192 and MS6193 [10] and appears to be in the same location in MS6194 and MS6203. We also identified a homolog of the MTase M.EcoST101V in MS6207 (100% nucleotide ID), however in MS6207 no methylation was observed at the corresponding 5’-CC^6m^**A**N_9_T<u>T</u>GG-3’ recognition site. Further investigation revealed that the specificity gene (*hsdS*) of RM.EcoST101V in the MS6207 genome was truncated at the 3’ end by the upstream insertion of a large ∼28 Kb composite transposon: Tn*6601*. This truncation resulted in the loss of DNA specificity and therefore loss of methylation and restriction activity. Additionally, in MS6207, the acquisition of this Tn*6601* resulted in the deletion of a 15.8 Kb genomic region immediately upstream of the IS*1R* (Figure 2). Tn*6601* is also present in both MS6203 and MS6201. In MS6203, Tn*6601* has also inserted upstream of the original IS*1R*, leaving the RD12 locus intact, however in MS6201, Tn*6601* has completely replaced the RD12 locus. In MS6204 however, Tn*6601* is not present and the absence of RM.EcoST101V is due to the absence of the RD12 locus, leaving only the conserved IS*1R* element remaining.

**Figure 2.**
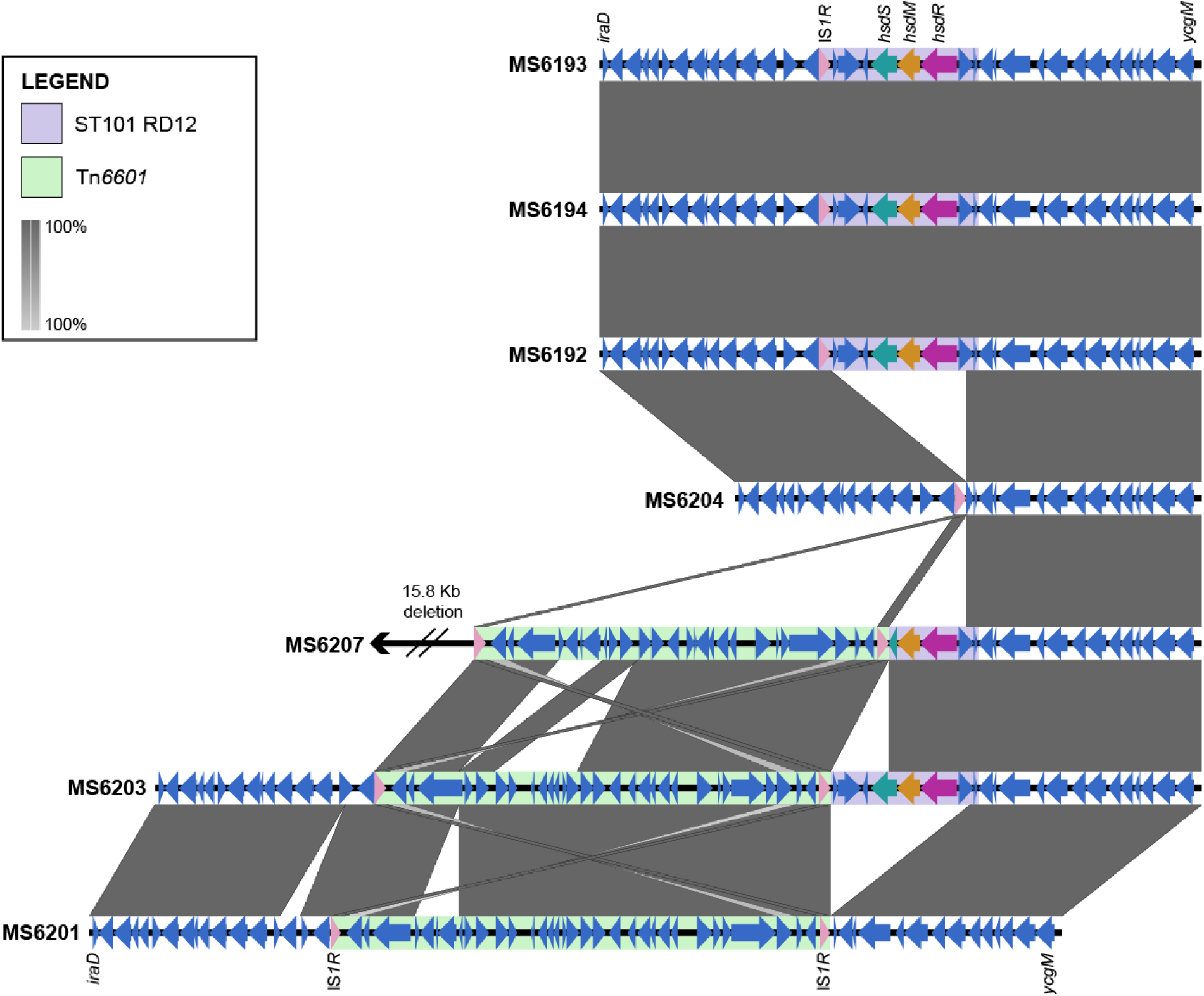
Conservation of RM.EcoST101V and the ST101 Region of Difference 12 (RD12) locus. Grey shading indicates nucleotide identity between sequences according to BLASTn (100%). MS6201 was reverse complemented for easier visualisation. Strains are in order as they appear in the ST101 phylogenetic tree [10]. ST101 Region of Difference (RD)12 (purple), IS*1R* flanked Tn*6601* (pale green), CDSs (blue), IS*1R* (pale-pink), *hsdS* specificity gene (teal), *hsdM* methyltransferase gene (orange), *hsdR* restriction gene (dark-pink). Image prepared using EasyFig.

To determine the distribution of Tn*6601* in ST101 strains, we included an additional seven published and publicly available Clade 1 ST101 draft genomes. All or most of this transposon is present in subclade 1.2 strains (MS6207, PI7, MS6203, NA086, NA084, MS6201 and NA099) (Supplementary Figure S2). This sequence of events is consistent with the acquisition of the IS*1R* flanked Tn*6601* into the same genomic locus as RD12 and then subsequent, independent transposition events leading to a) no change to the RD12 locus, b) truncation of the RD12 locus or c) loss of the RD12 locus. This result highlights how small recombination events at a single genomic locus between otherwise highly similar bacterial strains can result in global methylome changes within a single lineage.

#### Differential methylation due to acquisition of plasmid-borne RM.EcoST101IX

The 5’-CC^6m^**A**GN_6_R<u>T</u>TG-3’ recognition site encoded by the Type I RM system RM.EcoST101IX was also identified in the draft genomes of MS6194 and MS6204. In all four genomes that carry this Type I RM system (MS6192, MS6193, MS6194 and MS6204), this system is encoded on a pIP1206-like plasmid within the F-type plasmid backbone and is carried on the IS*1*-family flanked composite transposon Tn*6602* (Figure 3a). A BLAST comparison of RM.EcoST101IX confirmed that this Type I RM system is also present (100% nucleotide ID) in the original plasmid pIP1206 (Genbank accession: AM886293), with numerous homologs (>99% nucleotide ID) in other *E. coli, K. pneumoniae* and *Salmonella enterica* plasmid sequences (Supplementary Dataset, Table S5), highlighting the promiscuity of this RM system.

**Figure 3.**
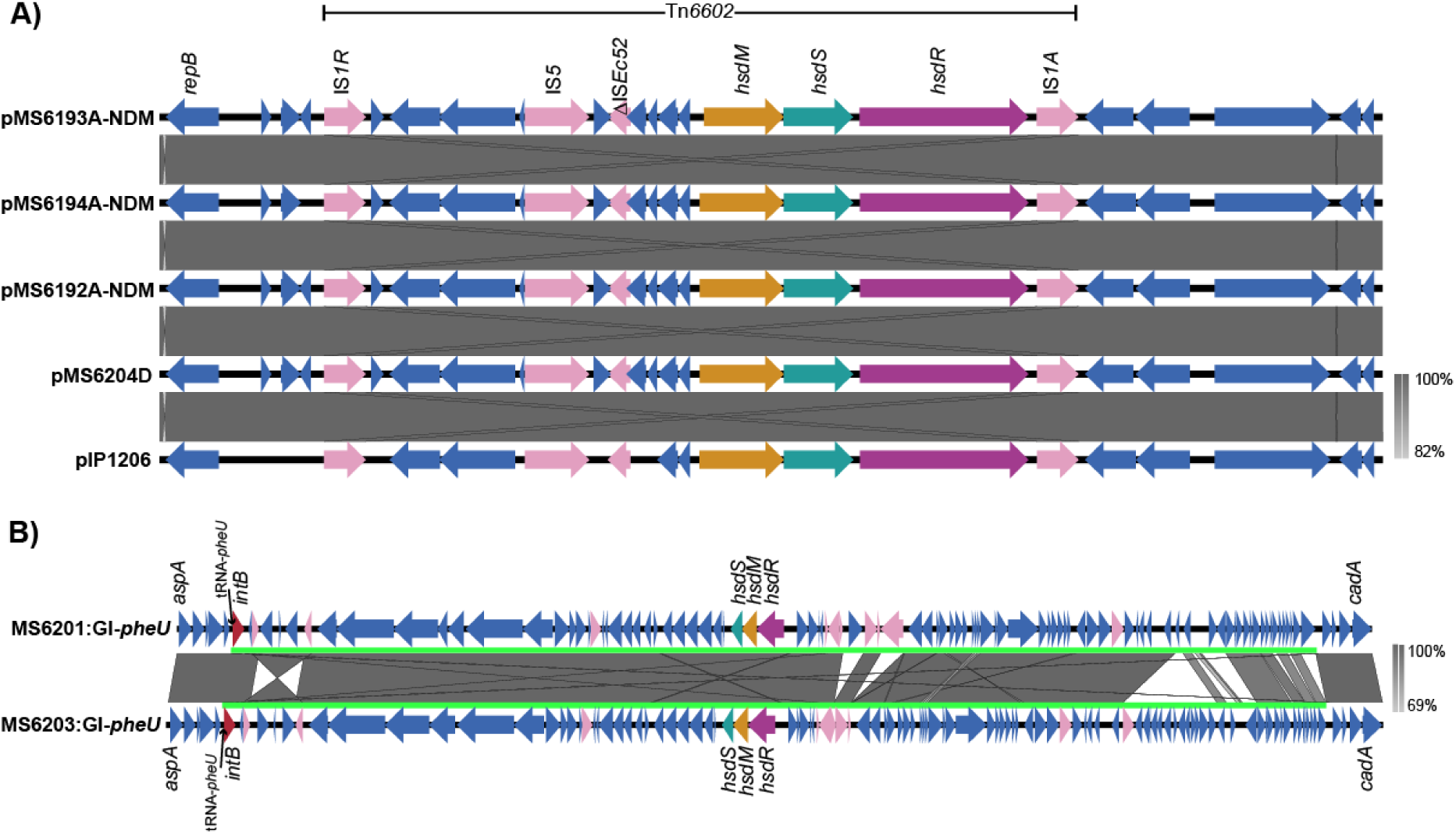
Genomic context of the ST101 Type I RM systems RM.EcoST101IX and RM.EcoST101XII. Schematic diagrams illustrating the genetic organisation and conservation of active RM systems. A) Tn*6602* encoded RM.EcoST101IX. B) Genomic Island GI-*pheU* encoded RM.EcoST101XII. Grey shading indicates nucleotide identity between sequences according to BLASTn. Key genomic features are indicated including integrase gene (red), Insertion sequences (pale-pink), *hsdS* specificity gene (teal), *hsdR* restriction gene (orange), *hsdM* methyltransferase gene (dark-pink), CDSs (pale blue). tRNA-*pheU* position labelled. GIs are indicated by the bright-green lines. Image prepared using EasyFig.

#### Differential methylation due to the acquisition of the chromosomal MGE-encoded RM.EcoST101XII

The methylated recognition site 5’-GC^6m^**A**N_5_G<u>T</u>TC-3’ is also characteristic of a Type I RM system and is present only in MS6201 and MS6203; it was not identified in MS6192 and MS6193. We searched the motif against REBASE and confirmed that it matches the recognition site of RM.Dso4321II, identified in the plant pathogen *Dickeya solani* D strain s0432-1 [31], a member of the Enterobacterales. Only a single Type I RM system was identified in MS6201 (designated RM.EcoST101XII), thus this is the probable cause of methylation at the 5’-GC^6m^**A**N_5_G<u>T</u>TC-3’ site. Comparisons of the specificity subunits S.Dso4321II and S.EcoST101XII however, indicate that they share only 81.22% amino acid identity, with several substitutions in the specificity domains (amino acids 4-183 and 244-366). While this recognition site has not previously been characterised in *E. coli*, the specificity gene is present in several *E. coli* genomes in REBASE (>99% amino acid identity) including one genome (*E. coli* O118:H16 str. 07-4255, Genbank accession: JASP01000001) that has associated PacBio sequence data. Despite the presence of this specificity gene in strain 07-4255, the recognition site was not identified in this genome. Further investigation revealed that the S and M subunits were present, however the R subunit was missing. It is currently unknown if the missing R subunit is the cause of the inactivity of this RM system in strain 07-4255.

To determine the genomic context of RM.EcoST101XII in MS6201, we characterised the surrounding genes. The presence of several IS elements, phage-like genes and hypothetical genes in close proximity to RM.EcoST101XII suggested carriage on a MGE. Comparative genomic analysis between our seven ST101 genomes characterised this region as a tRNA-*pheU* integrated GI (GI-*pheU*), different to the GI-*pheU* encoded in MS6192 and MS6193. MS6203 also contained a tRNA-*pheU* integrated GI, highly similar to that of the MS6201_GI-*pheU*, encoding the same Type I RM system RM.EcoST101XII as MS6201 (Figure 3b).

### DNA MTase distribution is reflected by differences in the accessory genome

To analyse the distribution of MTases across the ST101 lineage, we supplemented the seven PacBio sequenced genomes with 13 publicly available and published draft ST101 genomes and eight publicly available extraintestinal pathogenic *E. coli* (ExPEC) complete genomes (Supplementary Dataset, Table S2). *E. coli* ST101 genomes contain many MTases that are both conserved and variable across the lineage and other complete, reference ExPEC strains (Figure 4 and Supplementary Dataset, Table S6). M.EcoST101Dam, M.EcoST101Dcm and the YhdJ homolog M.EcoST101III are encoded in syntenic positions in all genomes, with all seven ST101 PacBio sequenced genomes showing methylation at the 5’-G^6m^**A**<u>T</u>C-3’ Dam recognition site. Aside from these core-genome conserved MTases, the distribution of all other ST101 MTases is consistent with the presence or absence of MGEs on which they are encoded (Supplementary Figure S3). For example, the Tn7-like Tn-encoded Type I RM system RM.EcoST101IV in MS6193 is present in only two other ST101 genomes (NA086 and NA084) that also carry the Tn7-like Tn and is completely absent in all other genomes surveyed. Likewise, the Type I RM system RM.EcoST101XII encoded on a tRNA-*pheU* integrated GI is present only in the two draft ST101 genomes MS6201 and MS6203. Other ST101 MTases however, show a variable distribution. For example, M.EcoST101I is encoded on Phi2 and shows a distribution consistent with the variability of this element in Clades 1 and 2 as well as the ExPEC complete genomes ED1a and CFT073. In MS6192, MS6193 and MS6194, M.EcoST101II is encoded on Phi6, however a homolog is also found in HVH 98 with several Phi6 gene modules also conserved in HVH 98. Interestingly, while M.EcoST101VI and M.EcoST101VII are both encoded on Phi7 in the complete genomes MS6192 and MS6193, their distribution differs throughout the ST101 lineage, which is likely due to differences in Phi7 gene content across the lineage.

**Figure 4.**
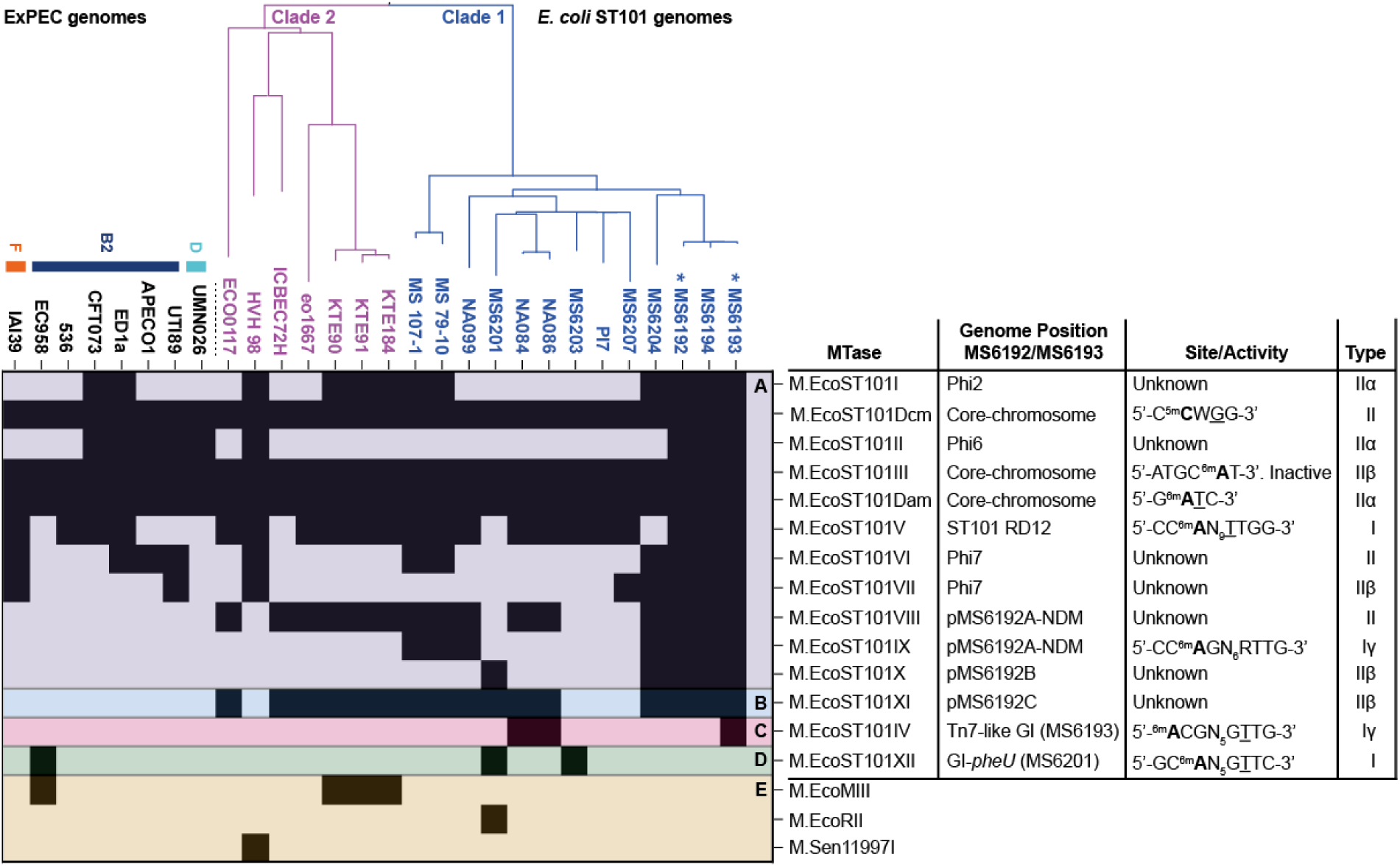
Distribution of MTases in the *E. coli* ST101 lineage. MTases conserved in *E. coli* MS6192 and MS6193 (A: purple), MTases encoded only in MS6192 (B: blue), MTases encoded only in MS6193 (C: pink), MTases encoded on the ST101 draft genomes only (D: green) and accessory ST101 MTases not encoded in *E. coli* MS6192 or MS6193 (E: orange) are shown along the X-axis. Strain identifiers are listed on the Y-axis, with ST101 strains ordered according to phylogenetic relationship. Black shading indicates a match of >=90% nucleotide identity with a minimum of 95% query coverage. Calculated by comparing the query sequences of *E. coli* MS6192 and MS6193 MTases and MTases defined as the gold standard from the REBASE database [14] to the complete genomes or draft assemblies of *E. coli* ST101 strains, as implemented in Seqfindr (http://github.com/mscook/seqfindr). Blastn results can also be found in Supplementary Dataset, Table S6.

Plasmid-borne MTases are also variably conserved throughout Clade 1 strains. The active, plasmid encoded Type I RM system RM.EcoST101IX is conserved across all Clade 1 strains that harbour Tn*6601*, carried on the pIP1206-like F-type plasmid. However, the Clade 1 isolates NA099, MS 79-10 and MS 107-1 also encode a homologous MTase. Further investigations reveal that NA099 shares an identical RM system to RM.EcoST101IX and contains an F-type plasmid, also encoding the *bla*_NDM-1_ locus. The MTase in MS 79-10 and MS 107-1 however shares only 97% nucleotide identity to the MTase M.EcoST101IX, with these genomes not encoding a *bla*_NDM-1_-positive F-type plasmid. Similarly, the Type II MTase M.EcoST101X is conserved across all Clade 1 strains that encode the pGUE-NDM-like FII plasmid, with an M.EcoST101X homolog (100% nucleotide ID) also present in MS6201. M.EcoST101X homologs are also present in several publicly available *E. coli* and *K. pneumoniae* plasmid sequences (Supplementary Dataset, Table S7), suggesting that despite inactivity under normal laboratory growth conditions, this MTase may have an important biological function. Lastly, homologs of M.EcoST101XI encoded on the IncI1 plasmid pMS6192C are present in the majority of ST101 strains, even in genomes that do not carry IncI1 plasmids.

Using the REBASE Gold Standard database (MTases that have been experimentally validated) and removing redundancy, we identified three additional accessory MTase genes that were not encoded within the complete genomes MS6192 and MS6193 (Figure 4). One of these MTases, M.EcoRII is part of a Type II RM system, present on an FII plasmid in MS6201 and is a predicted Dcm homolog. The Clade 2 isolates KTE184, KTE91 and KTE90 contain a Type I MTase similar (96% nucleotide ID) to M.EcoMIII from the ExPEC complete genome EC958 [32]. Lastly, the Clade 2 strain HVH 98 also encodes a Type I MTase, homologous (92% nucleotide ID) to M.Sen11997I from the *Salmonella enterica* subsp. enterica serovar Chester strain ATCC 11997 [33].

## Discussion

We have previously characterised the role of MGEs in the carriage of *bla*_NDM-1_, conferring carbapenem resistance in the two *E. coli* ST101 complete genomes (MS6192 and MS6193) and five draft genomes (MS6194, MS6201, MS6203, MS6204 and MS6207) [10]. In the present study, we used these genomes and the kinetics of PacBio SMRT sequencing to bioinformatically characterise DNA MTases, assign recognition sites with their cognate MTase and to define the genomic context of MTases within our collection, facilitating the first comprehensive methylome analysis of the ST101 lineage. We identified 14 DNA MTases and eight ^6m^A recognition sites, including one novel site that could not be assigned to its cognate MTase. We also showed that eight MTases shared by MS6192 and MS6193 were either inactive under the growth conditions tested or responsible for ^5m^C methylation, which was not characterised in this study. Transcriptionally silent MTases may be active under specific circumstances such as stress induction or changes in environment. It is possible that cloning and expression of these genes via a plasmid system in a MTase-free strain of *E. coli*, as has been performed previously for other MTases [34], could reveal their target specificity. Overall, our capacity to resolve complex MGEs and define the genomic context of MTases within the ST101 lineage has revealed strain, clade and lineage-wide methylome heterogeneity.

There is an almost identical methylation profile between the two complete ST101 genomes MS6192 and MS6193, however we show that the acquisition of a single, active RM system (RM.EcoST101IV) encoded on the Tn7-like Tn (present only in MS6193) resulted in 788 differentially methylated sites. While more than 96% of sites were within intragenic regions of the genome, 27 sites were within intergenic regions, with all but one located in putative promoter regions (which we defined as ≤300 bp from a start codon). Thus, it is possible that methylation of the RM.EcoST101IV site 5’-^6m^**A**CGN_5_G<u>T</u>TG-3’ could result in an indirect role in gene regulation. While the gene regulatory role of orphan MTases such as Dam has previously been demonstrated [11], there are also examples of acquisitions of RM systems that have caused differential methylation patterns and thus differential gene regulation. For example, comparisons of the knockout mutant of the Type II RM system RM.EcoGIII encoded on the Shiga toxin phage, to the wild-type *E. coli C227*-11 strain led to more than a third of all genes differentially expressed [34], indicating that acquired MTases encoded on MGEs can result in significant changes to gene expression. Future work will involve analysing the intersection of the methylome and transcriptome via RNA sequencing methods.

Currently, it is unknown whether the additional RM system RM.EcoST101IV could generate barriers of DNA exchange and influence the gene pool available to MS6193 however, RM systems do have a role in maintaining species identity and restricting horizontal gene transfer in some species. For example, in *Neisseria meningitidis*, the distribution of RM systems is consistent with its phylogenetic clade structure [35]. Intraclade HGT was significantly more likely than interclade HGT, highlighting that RM systems generate barriers to DNA exchange and are involved in the evolution of distinct lineages [35]. In *Staphylococcus aureus*, a mutation in the restriction subunit (*hsdR*) of the Type I RM system Sau1 is vital for plasmid transformation of the laboratory strain *S. aureus* RN4220, allowing uptake of foreign DNA [36]. Additionally, distinct variants of two specificity units (*hsdS*) encoded on GIs were identified across the different lineages of S. aureus, indicating lineage-specific sequence specificity [36]. In *Burkholderia pseudomallei*, each lineage contained a distinct complement of RM systems, which caused clade-specific methylation patterns. Transformation with reporter plasmids that carried specific restriction sites impeded the ability of the *E. coli* strains encoding distinct *B. pseudomallei* RM systems to be transformed [37]. It is therefore predicted that these lineage-specific RM systems partition the species by restricting HGT and inhibiting uptake of non-self-DNA [37]. Whether RM systems within ST101 present a significant barrier to HGT between lineages has yet to be elucidated and represents an area of future research interest.

In the seven PacBio sequenced genomes, we characterised only a single ST101 MTase capable of ^5m^C methylation (*dcm* encoded by M.EcoST101Dcm, which methylates 5’-C^5m^**C**W<u>G</u>G-3’ sites) that has previously been characterised in *E. coli* [38]. However, our ability to detect ^m5^C methylation was limited. The kinetics of ^5m^C methylation are subtle and spread over several bases as the modification is hidden in the major groove of the DNA, limiting the effectiveness of the detection algorithm [20]. This could have been overcome by increasing the number of SMRT cells used and thus throughput, increasing the sequencing coverage to 250X [22]. Alternatively, enzymatic conversion via Ten-eleven translocation (Tet) treatment to convert ^5m^C to 5-carboxylcytosine increases the size of the modification, enhancing the kinetic signal [39]. However, this conversion is sometimes incomplete and even with complete conversion, ^5m^C isn’t always detected at complete levels [40]. Thus, we focused our study on the dominant ^6m^A modifications in *E. coli*.

To date, eleven *E. coli* methylomes have been published, including the *E. coli* strains DH5α, BL21(DE3) and Bal225 [41], C277-11 [34], RM13514 and RM13516 [42], EC958 [32], CFT073 and K-12 substr. MG1655 [23] and 95NR1 and 95JB1 [43]. These studies have highlighted the diversity of MTases across *E. coli*, with the MTase complement and site specificities varying significantly even between members of the same phylogroup and ST. However, in general, each study was restricted to a very small number of genomes, limiting our knowledge of MTase conservation across whole lineages. Currently, this is only the second study of the distribution of MTases within strains of an *E. coli* lineage, where we first noted the importance of MGEs in the distribution of MTases and showed lineage-specific methyltransferase patterns in the UPEC ST131 clone [32]. By characterising the genomic context of all MTases in our two ST101 complete genomes and active MTases in our five draft genomes, we showed that the majority are encoded on MGEs. Including an additional 13 published and publicly available draft genomes confirmed that variation in MTases within the ST101 lineage was mostly due to MGE differences between genomes.

Furthermore, there were limited numbers of accessory MTases identified that were not encoded within either MS6192 or MS6193. While our identification of accessory MTases was restricted as we only compared against MTases that have been experimentally shown to possess methylation activity (REBASE Gold Standard) [14], these limited numbers of accessory MTases may indicate that MTases act as a barrier to HGT within the lineage.

Our analysis of the ST101 methyome shows that even within a single clade, substantial differences in MTase content can occur, highlighting the need for multiple PacBio genomes across all clades to reveal the full extent of epigenomic diversity within a lineage. Additionally, our findings demonstrate the significant role of MGEs in enabling very similar bacterial strains to rapidly acquire genome-wide differences in their methylome, highlighting the expanding role of MGEs in *E. coli* evolution. Further work studying the intersection between the methylome and transcriptome will expand our understanding of the functional roles of DNA methylation in bacteria and provide new insights into how strain and lineage-specific methylome changes drive host adaptation.

## Authors contributions

Conceptualisation: MMA, BMF, MAS and SAB. Investigation: MMA, BMF. Formal Analysis: MMA. Visualisation: MMA. MDP, KMP and DLP assisted in clinical/wet-lab experiments. BMF and LWR assisted in data analysis. KGC, TMC and WFY performed the PacBio sequencing. Resources: TRW, KGC, MAS and SAB. Supervision: BMF, MAS and SAB. Writing (Original Draft Preparation): MMA, BMF and SAB. Writing (Review and Editing): MMA, MDP, KGC, MAS and SAB. All authors contributed to the final review and edits.

## Funding

This work was supported by grants from the Australian National Health and Medical Research Council (G1033799) and from the University of Malaya High Impact Research (HIR) Grants (UM-MOHE HIR Grant UM.C/625/1/HIR/MOHE/CHAN/14/1, Grant H-50001-A000027 and FP022-2018A). MAS is supported by a NHMRC Senior Research Fellowship (G1106930). SAB is supported by a NHMRC Career Development Fellowship (G1090456). MMA and LWR were supported by an Australian Government Research Training Program Scholarship. The funding bodies had no role in the design of the study and the collection, analysis, and interpretation of data or in writing of the manuscript.

## Conflicts of interest

The authors declare that there are no conflicts of interest.

## Notes

### Competing Interest Statement

The authors have declared no competing interest.

