## Supplementary Appendix for "Strain and lineage-level methylome heterogeneity in the multi-drug resistant pathogenic *Escherichia coli* ST101 clone"

**Supplementary Figures.**

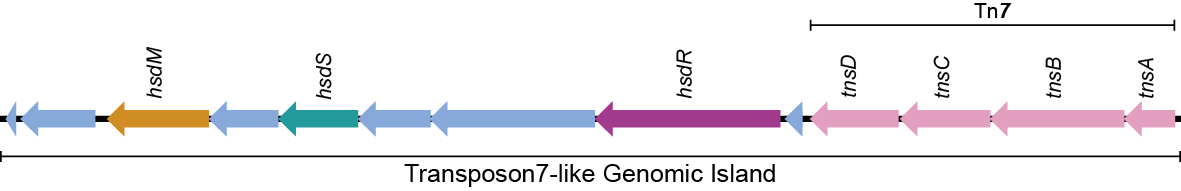

**Figure S1. Acquisition of the Transposon *7*-like Transposon encoding RM.EcoST101VIII in MS6193.** Methyltransferase gene *hsdM* (orange), specificity gene *hsdS* (teal) and restriction gene *hsdR* (dark-pink). Genes associated with transposition (light-pink); other CDSs (blue). Image prepared using EasyFig [1].

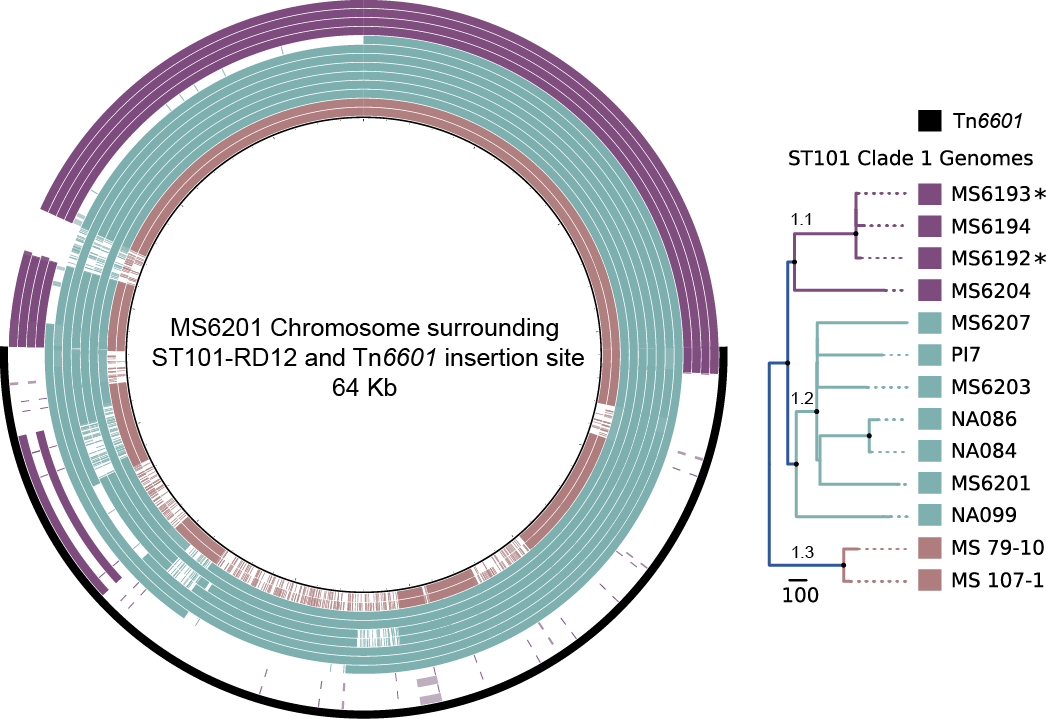

**Figure S2. Genomic map highlighting the ST101 IS*1R* flanked Resistance Island Tn*6601*.** Visualisation of the *E. coli* MS6201 genome surrounding the IS*1R* flanked composite transposon Tn*6601* compared to 12 *E. coli* ST101 Clade 1 genomes. The degree of coloured shading indicates nucleotide identity between MS6201 and each *E. coli* ST101 genome. Nucleotide comparisons are coloured based on identity between 70 and 100% (dark shading = high identity, light shading = low identity). *E. coli* Clade 1 ST101 genomes are arranged (right) according to their previously defined phylogenetic relationship [2] as follows from the innermost ring: Subclade 1.3 (brown): MS 107-1 and MS 79-10; Subclade 1.2 (green): NA099, MS6201, NA084, NA086, MS6203, PI7, MS6207; Subclade 1.1 (purple): MS6204, MS6192, MS6194, MS6193. *Denotes complete ST101 genomes. Outer ring indicates position of Tn*6601* (black). Image prepared using BRIG [3].

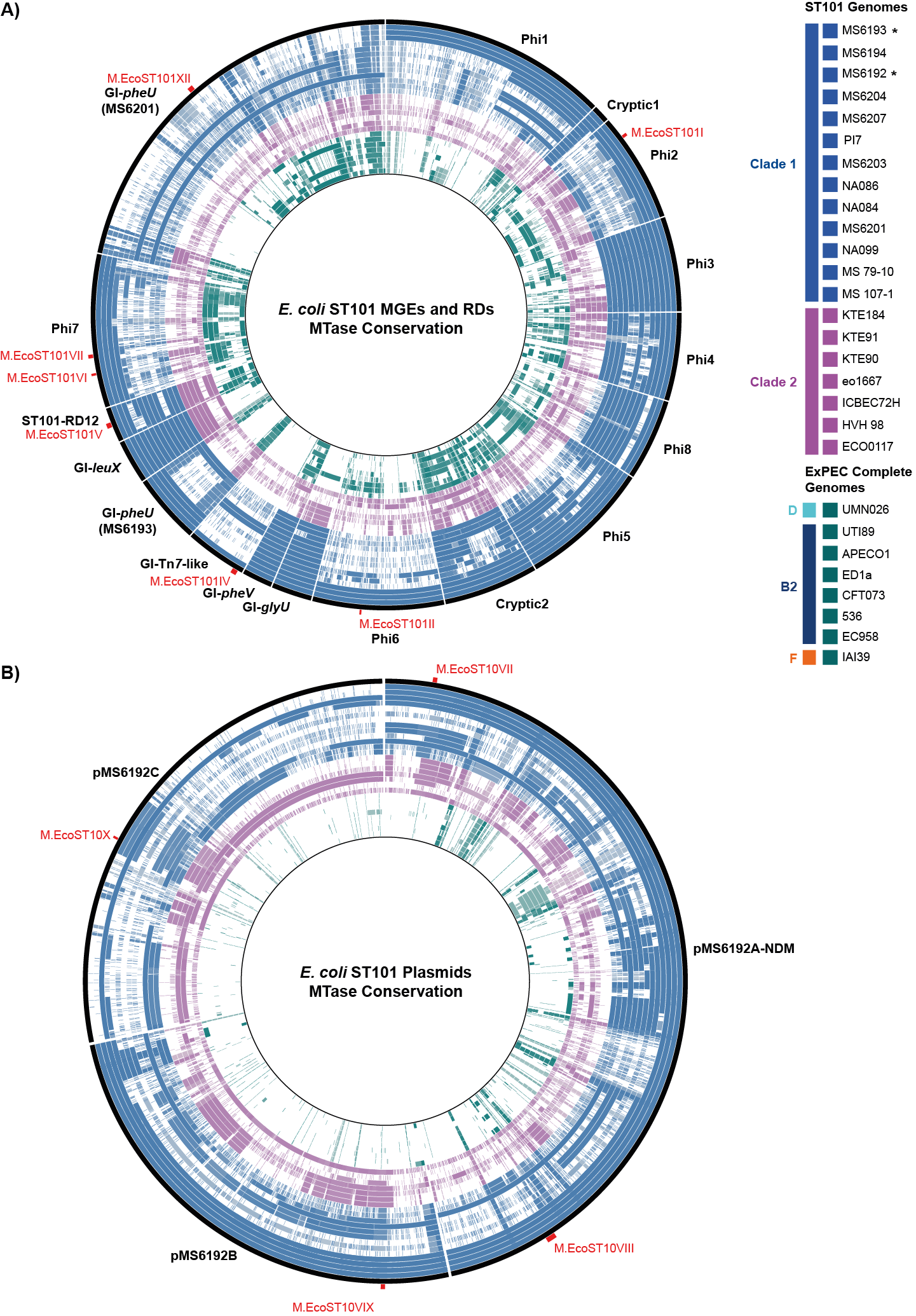

**Figure S3. Distribution of mobile genetic elements and DNA methyltransferases in *E. coli* ST101 and representative ExPEC genomes.** The degree of coloured shading indicates the nucleotide identity between each genome. A) Chromosomal MGEs and RDs B) Plasmids. Nucleotide comparisons are coloured based on identity between 70-100% (light-dark shading). *E. coli* ST101 genomes are arranged according to their previously defined phylogenetic relationship [2] as follows from the outermost ring: Clade 1 (blue): MS6193*, MS6194, MS6192*, MS6204, MS6207, PI7, MS6203, NA086, NA084, MS6201, NA099, MS 79-10 and MS107-1; Clade 2 (pink): KTE184, KTE91, KTE90, eo1667, ICBEC72H, HVH 98 and ECO0117. ExPEC representative complete genomes (green) are arranged in order: Phylogroup D: UMN026, Phylogroup B2: UTI89, APECO1, ED1a, CFT073, 536, EC958 and Phylogroup F: IAI39. *Denotes complete ST101 genomes. Black outermost ring indicates reference MGE or plasmid (labelled in black). Accessory ST101 MTases indicated in Red. Image prepared using BRIG [3].
